# Sensitive period for cognitive repurposing of human visual cortex

**DOI:** 10.1101/402321

**Authors:** Shipra Kanjlia, Rashi Pant, Marina Bedny

## Abstract

Studies of sensory loss are a model for understanding the functional flexibility of human cortex. In congenital blindness, subsets of visual cortex are recruited during higher-cognitive tasks, such as language and math tasks. Is such dramatic functional repurposing possible throughout the lifespan or restricted to sensitive periods in development? We compared visual cortex function in individuals who lost their vision as adults (after age 17) to congenitally blind and sighted blindfolded adults. Participants took part in resting-state and task-based fMRI scans during which they solved math equations of varying difficulty and judged the meanings of sentences. Blindness at any age caused “visual” cortices to synchronize with specific fronto-parietal networks at rest. However, in task-based data, visual cortices showed regional specialization for math and language and load-dependent activity only in congenital blindness. Thus, despite the presence of long-range functional connectivity, cognitive repurposing of human cortex is limited by sensitive periods.

## Introduction

Studies of sensory loss provide a model for understanding cortical flexibility (Bavelier and Neville 2002). In arm amputees, the hand area of somatosensory cortex responds to stimulation of the face (Pons, Garraghty, Ommaya, Kaas, 1991). The auditory cortices of deaf individuals respond to visual stimuli and the visual cortices of blind individuals respond to sound and touch, a phenomenon termed cross-modal plasticity (Sadato et al. 1996; Cohen et al. 1997; Büchel et al. 1998; Bavelier and Neville 2002; Collignon et al. 2011; Watkins et al. 2013; Almeida et al. 2015). Even in such reorganization, cortex typically retains elements of its original functions. For example, in arm amputation, somatosensory regions corresponding to the hand continue to perform somatosensation, but now over input from the face (Pons, Garraghty, Ommaya, Kaas, 1991). Prior work with blind individuals also suggests that some “visual” cortex functions preserved in congenital blindness (Striem-Amit, Cohen, et al. 2012; Striem-Amit, Dakwar, et al. 2012; Striem-Amit and Amedi 2014).

However, a growing body of evidence suggests that, in congenital blindness, visual cortices take on entirely different functions from their typical role in visual perception. In addition to non-visual sensory responses, many of the cross-modal responses observed in visual cortex of blind individuals appear to reflect higher-cognitive operations, such as language and mathematical processing (for review see Bedny, 2017). Parts of the visual cortex, such as lateral occipital and ventral occipito-temporal regions, are active during sentence comprehension and increase activity with the grammatical complexity of spoken sentences (Bedny et al., 2011; Lane et al., 2015). A separate dorsal visual region (right middle occipital gyrus, rMOG) is active during math calculation and increases activity with the difficulty of math equations (Kanjlia et al. 2016; Amalric et al. 2017). There is evidence that visual cortex activity during higher-cognitive tasks in blindness is behaviorally relevant. TMS to occipital cortex causes congenitally blind but not sighted individuals to make errors when reading Braille and when generating semantically appropriate verbs to heard nouns (Cohen et al. 1999; Amedi et al. 2003, 2004; Merabet et al. 2004). Even at rest, activity in visual cortices is synchronized with higher-cognitive fronto-parietal networks in congenitally blind but not sighted individuals (Liu et al. 2007; Bedny et al. 2011; Watkins et al. 2012; Kanjlia et al. 2016). Recruitment of visual cortices for higher-cognitive functions is the most extreme example of cortical cognitive repurposing identified to date, since language and mathematics are cognitively and evolutionarily distant from low-level vision.

What are the limits on such cognitive reorganization in cortex? Does human cortex retain the ability to support a wide range of cognitive functions throughout the lifespan? Alternatively, is such drastic functional repurposing uniquely possible during sensitive periods of development?

It is generally established that plasticity in the developing brain is enhanced relative to the mature brain. The most well studied example of this phenomenon comes from monocular visual deprivation. When one eye does not receive typical input during a critical period in development, visual cortex neurons that would normally respond to the deprived eye are overtaken by input from the dominant or “good” eye (Hubel and Wiesel 1970). Analogously in humans, dense cataracts in one eye during the first years of life but not afterwards cause impairments in visual acuity, even after the cataract is removed (Banks et al. 1975; Lewis and Maurer 2005). Recent research in the mouse model has uncovered local-circuit neurophysiological mechanisms that regulate sensitive period plasticity and distinguish it from other forms of learning. Sensitive period opening and closure involves shifts in the excitatory/inhibitory balance and the closure of sensitive periods coincides with formation of perineuronal nets, which dampens synaptic plasticity (Pizzorusso, 2002; Hensch, 2005; Bavelier *et al.*, 2010). Thus, local circuit plasticity during sensitive periods is mediated by specific neurophysiological mechanisms.

Whether the capacity of cortex to take on novel cognitive functions similarly depends on sensitive period plasticity remains unknown. As noted above, some functional plasticity is possible, even in adulthood (Merzenich et al. 1983, 1984; Kaas 1991). For example, amputation of a limb causes neighboring cortical representations of intact body parts to expand into deaffrented somatosensory cortices (Calford and Tweedale 1988; Pascual-Leone et al. 1996, 2005; Borsook et al. 1998; Röricht et al. 1999). This activation appears to be functionally relevant as TMS to the newly deaffrented arm region of somatosensory cortex induces sensations in the face and biceps (Pascual-Leone et al. 1996; Röricht et al. 1999). Arguably, however, the functional plasticity observed in amputation is relatively subtle, as compared to that seen in blindness or deafness. Is more dramatic functional repurposing of cortex circumscribed to sensitive periods of development?

Some evidence for the idea that visual cortices assume different functions in congenital and adult-onset blindness comes from studies of auditory motion and spatial perception. Dorsal visual areas that preferentially respond to sound localization in congenital blindness do not show such cross-modal recruitment in adult-onset blindness (Haxby et al. 1991; Goodale and Milner 1992; Voss et al. 2006; Collignon et al. 2013a). Visual motion processing area, MT, only shows enhanced auditory motion processing in individuals who lose their vision early in life, not later in life (Jiang et al. 2016). Such evidence suggests that the capacity of cortex to take on novel functions in adulthood is restricted.

However, studies of higher-cognitive plasticity in visual cortex of adult-onset blind individuals have thus far yielded mixed results. Consistent with the idea of sensitive periods, one study reported that V1 responds more to sentences than non-verbal sounds only in those who are congenitally blind (Bedny et al. 2012). On the other hand, even in adult-onset blindness, visual cortices appear to be active during higher-cognitive tasks, such as Braille reading, phonological judgments of spoken words and sentence comprehension, although it is not yet clear what such activity reflects (Cohen et al. 1999; Burton and McLaren 2006; Burton et al. 2011). A recent study also found that resting-state activity of visual cortices becomes synchronized with that of Broca’s area in adult-onset blindness, suggesting repurposing of visual cortices for language even in adulthood (Sabbah et al. 2016).

None of previous studies, however, directly address the question of whether visual cortices are sensitive to higher-cognitive information in adult-onset blindness. The most compelling evidence for visual cortex involvement in higher-cognitive functions in congenital blindness comes from studies that manipulate fine-grained higher-cognitive information, such as the grammatical complexity of sentences and difficulty of math equations (Röder et al. 2002; Bedny et al. 2011; Lane et al. 2015; Kanjlia et al. 2016). By contrast, all prior work with adult-onset blind individuals has compared higher-cognitive tasks to a resting baseline or low-level perceptual control condition, making it difficult to determine what cognitive processes visual cortex activity truly reflects in the adult-onset blind population (Cohen et al. 1999; Burton and McLaren 2006; Burton et al. 2011). If the extreme cognitive flexibility of cortex is restricted to a sensitive period, visual cortices of adult-onset blind individuals should not respond to manipulations of higher-cognitive information.

A further open question concerns whether cognitive repurposing, as measured by task-based responses, follows a similar developmental time-course as changes in resting-state connectivity. As noted above, in congenital blindness, resting-state activity in visual cortices becomes synchronized with that of fronto-parietal higher-cognitive networks. These resting-state changes are region and network-specific. “Visual” regions that are active during mathematical processing show correlated activity with fronto-parietal number networks, even at rest, whereas those that respond to grammatical and semantic information during language tasks are correlated with Broca’s area (Bedny et al. 2009; Kanjlia et al. 2016). It is not known whether such region-specific increases in functional connectivity of visual cortex follow a sensitive period and, if so, whether this sensitive period aligns with that of task-based responses. Answering this question could provide general insights into the relationship between task-based and resting-state connectivity measures.

In the current study, we addressed these open questions by comparing task-based activation and resting-state functional connectivity across adult-onset blind (blind after 17-years-of-age), congenitally blind and blindfolded sighted participants. First, we asked whether visual cortices of adult-onset blind individuals show regional specialization for math as opposed to language and whether they show load dependent responses during higher-cognitive tasks--in particular, during symbolic mathematical reasoning. As noted above, in congenitally blind individuals, a dorsal visual area (rMOG) is more active during symbolic a math task (e.g. 27-12=x) than during a matched sentence comprehension task and activity in the rMOG increases with the difficulty of math equations (Kanjlia et al. 2016). Here we tested whether the rMOG of adult-onset blind individuals has a similar functional profile. Second, we tested whether adult-onset blind individuals, like the congenitally blind group, show higher resting-state functional connectivity between the rMOG and the fronto-parietal number network and higher functional connectivity between a language-responsive visual cortex area (ventral occipito-temporal cortex or VOT) and prefrontal language areas.

## Materials and Methods

### Participants

Nineteen blind-folded sighted (age=21.45-75.49 years, mean=45.61, SD=16.03; 9 female), 13 adult-onset blind (age=34.74-74.72, mean=57.18, SD=11.77; 3 female) and 20 congenitally blind (age=19.34-70.12, mean age=46.08 years, SD=16.80; 15 female) participants contributed data to the current study (Table 1). Seven additional participants were scanned but excluded from all analyses because overall accuracy on the math and language tasks fell below 60% (5 congenitally blind) or because of incomplete coverage of the occipital lobe (2 sighted).

**Table 1.**
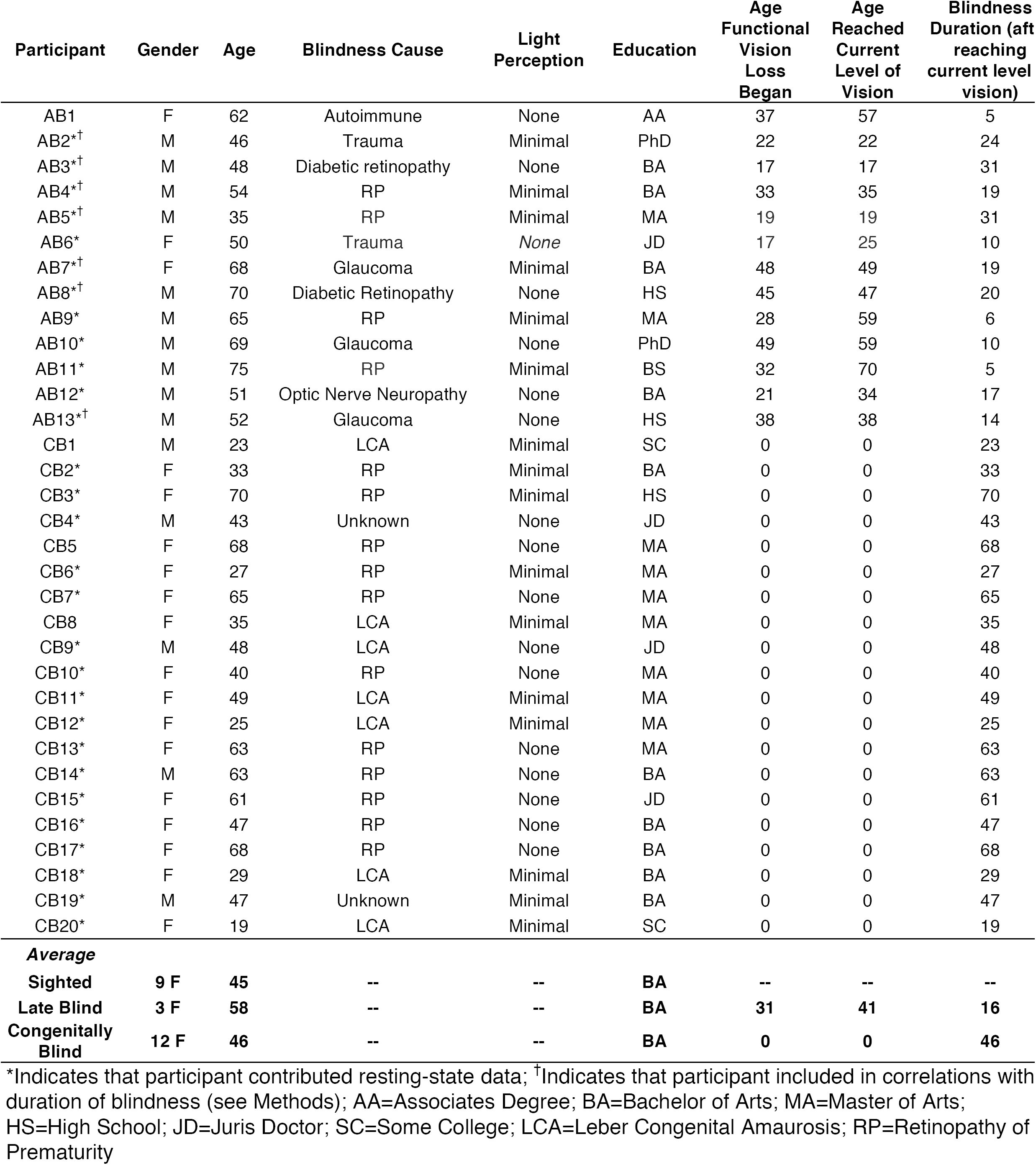
Demographic information for participants in math task.

| Participant | Gender | Age | Blindness Cause | Light Perception | Education | Age Functional Vision Loss Began | Age Reached Current Level of Vision | Blindness Duration (aft reaching current level vision) |
| --- | --- | --- | --- | --- | --- | --- | --- | --- |
| AB1 | F | 62 | Autoimmune | None | AA | 37 | 57 | 5 |
| AB2*<sup>†</sup> | M | 46 | Trauma | Minimal | PhD | 22 | 22 | 24 |
| AB3*<sup>†</sup> | M | 48 | Diabetic retinopathy | None | BA | 17 | 17 | 31 |
| AB4*<sup>†</sup> | M | 54 | RP | Minimal | BA | 33 | 35 | 19 |
| AB5*<sup>†</sup> | M | 35 | RP | Minimal | MA | 19 | 19 | 31 |
| AB6* | F | 50 | Trauma | None | JD | 17 | 25 | 10 |
| AB7*<sup>†</sup> | F | 68 | Glaucoma | Minimal | BA | 48 | 49 | 19 |
| AB8*<sup>†</sup> | M | 70 | Diabetic Retinopathy | None | HS | 45 | 47 | 20 |
| AB9* | M | 65 | RP | Minimal | MA | 28 | 59 | 6 |
| AB10* | M | 69 | Glaucoma | None | PhD | 49 | 59 | 10 |
| AB11* | M | 75 | RP | Minimal | BS | 32 | 70 | 5 |
| AB12* | M | 51 | Optic Nerve Neuropathy | None | BA | 21 | 34 | 17 |
| AB13*<sup>†</sup> | M | 52 | Glaucoma | None | HS | 38 | 38 | 14 |
| CB1 | M | 23 | LCA | Minimal | SC | 0 | 0 | 23 |
| CB2* | F | 33 | RP | Minimal | BA | 0 | 0 | 33 |
| CB3* | F | 70 | RP | Minimal | HS | 0 | 0 | 70 |
| CB4* | M | 43 | Unknown | None | JD | 0 | 0 | 43 |
| CB5 | F | 68 | RP | None | MA | 0 | 0 | 68 |
| CB6* | F | 27 | RP | Minimal | MA | 0 | 0 | 27 |
| CB7* | F | 65 | RP | None | MA | 0 | 0 | 65 |
| CB8 | F | 35 | LCA | Minimal | MA | 0 | 0 | 35 |
| CB9* | M | 48 | LCA | None | JD | 0 | 0 | 48 |
| CB10* | F | 40 | RP | None | MA | 0 | 0 | 40 |
| CB11* | F | 49 | LCA | Minimal | MA | 0 | 0 | 49 |
| CB12* | F | 25 | LCA | Minimal | MA | 0 | 0 | 25 |
| CB13* | F | 63 | RP | None | MA | 0 | 0 | 63 |
| CB14* | M | 63 | RP | None | BA | 0 | 0 | 63 |
| CB15* | F | 61 | RP | None | JD | 0 | 0 | 61 |
| CB16* | F | 47 | RP | None | BA | 0 | 0 | 47 |
| CB17* | F | 68 | RP | None | BA | 0 | 0 | 68 |
| CB18* | F | 29 | LCA | Minimal | BA | 0 | 0 | 29 |
| CB19* | M | 47 | Unknown | Minimal | BA | 0 | 0 | 47 |
| CB20* | F | 19 | LCA | Minimal | SC | 0 | 0 | 19 |
| **Average** |  |  |  |  |  |  |  |  |
| **Sighted** | **9 F** | **45** | -- | -- | **BA** | -- | -- | -- |
| **Late Blind** | **3 F** | **58** | -- | -- | **BA** | **31** | **41** | **16** |
| **Congenitally Blind** | **12 F** | **46** | -- | -- | **BA** | **0** | **0** | **46** |

\*Indicates that participant contributed resting-state data; <sup>†</sup>Indicates that participant included in correlations with duration of blindness (see Methods); AA=Associates Degree; BA=Bachelor of Arts; MA=Master of Arts; HS=High School; JD=Juris Doctor; SC=Some College; LCA=Leber Congenital Amaurosis; RP=Retinopathy of Prematurity

All blind participants had at most minimal light perception at the time of the experiment and had lost their vision due to pathology at or anterior to the optic chiasm and not due to brain damage. All participants reported having no cognitive or neurological disabilities. Participants with adult-onset blindness became blind (reached their current level of vision) after the age of 17 (mean=40.85, SD=17.36, min=17, max=70) and were blind for an average of 16.11 years after reaching their current level of vision (SD=8.99, min=4.72, max=31.35) (Table 1).

Forty-three blind-folded sighted (25 female; mean age=34.12 years, SD=14.33, min=18.88, max=63.19), 12 adult-onset blind (2 female; mean age=56.79, SD=12.21, min=34.74, max=74.75) and 25 (18 female; mean age=46.63, SD=16.91, min=18.81, max=72.98) congenitally blind individuals contributed resting-state data. A subset of participants who contributed resting-state data also participated in the task-based fMRI experiment (indicated with asterisk in Table 1).

Task data from all 19 sighted participants and 16 congenitally blind participants as well as resting-state data from 9 sighted and 12 congenitally blind were previously published (Kanjlia et al. 2016).

### Behavioral Task

Participants performed auditory math and language-control tasks while undergoing fMRI. Stimuli were presented in American English and were delivered to the participant through MRI compatible headphones. On math trials, participants heard two math equations each containing an unknown variable (e.g. 7-2=x). Equations lasted 3.5 seconds each and were separated by a 2.75 second delay. Participants pressed one of two buttons to indicate whether the value of x in the two equations was the same (4 seconds to respond). Participants were able to respond at any point after the onset of the second math equation or sentence, thus response times in Fig. 1 are relative to the onset of the second stimulus.

**Figure 1.**
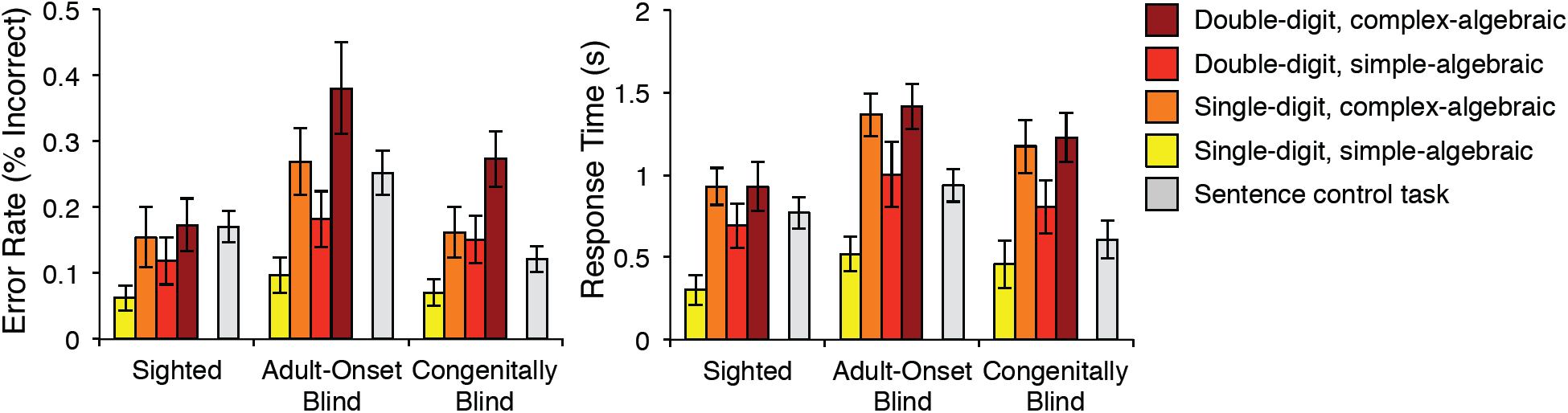
Behavioral Performance. Error rates (left) and response times (relative to offset of second stimulus; right) for all conditions in math task (warm colors) and language control task (grey). Error bars show standard error of the mean.

The format of language trials was identical to that of math trials except participants heard 2 sentences and indicated whether the meaning of the two sentences was the same. One of the sentences was always in active voice and the other was in passive voice. On “different” trials, who-did-what-to-whom was switched from one sentence to the other while all nouns and verbs remained identical. Half of the language trials had an object relative construction and half had a subject relative construction (two total language conditions). These two language conditions were not compared in this study.

The difficulty of math equations was varied using two orthogonal manipulations (four total math conditions). Half of the equations contained all single-digit numbers (e.g. 7-2=x) and half contained all double-digit numbers (e.g. 27-12=x). Orthogonally, in half of the equations, the unknown variable x was isolated on the right side of the equation (algebraically simple; e.g. 7-2=x), while the other half required manipulation to isolate x (algebraically complex; e.g. x-2=7). Double-digit math equations never required “carry-over” to reach a solution, thus reducing any differences in working memory demands across the double- and single-digit conditions. By contrast, the algebraic complexity manipulation may tax both numerical and working memory processes (Maruyama et al. 2012; Monti et al. 2012).

Each pair of math equations and sentences was presented once throughout the experiment. The experiment was divided into 6 runs each with 24 trials (16 math trials and 8 language trials). The 4 math conditions and 2 language conditions (6 total conditions) were counterbalanced in a Latin square design across all 6 runs. A small number of participants completed fewer than 6 runs of the experiment (4 AB, 2 CB, and 7 S completed 5 runs and 2 S completed 4 runs). Thus, there was a total of 96 unique math trials and 48 unique language trials in the experiment.

All participants (including adult-onset blind and congenitally blind participants) were blind-folded throughout the experiment and resting-state scans.

### MRI Data acquisition

A 3T Phillips scanner was used to collect whole-brain MRI anatomical and functional data. T1-weighted anatomical images were collected in 150 1-mm axial slices (1-mm isotropic voxels). Functional BOLD data were collected in 36 3-mm axial slices (2.4 x.4 x 3mm voxels; TR=2 seconds, TE=0.03 seconds; FOV xyz=171.79 x 192 x 107.5 mm). The same functional parameters were used to collect 1-4 8-minute resting-state scans during which participants were instructed to rest and remain awake.

### fMRI Data analysis

fMRI Data were analyzed using Freesurfer, FSL, HCP workbench and custom in-house software. Data were motion corrected, high-pass filtered (128 seconds), mapped to the cortical surface using the standard Freesurfer pipeline, spatially smoothed on the surface (6-mm FWHM Gaussian kernel), and prewhitened to remove temporal autocorrelation.

Task-based fMRI data were analyzed using a standard general linear model (GLM). Each of the four math conditions and each of the 2 language conditions were entered as predictors in the GLM after convolving with the canonical hemodynamic response function. First temporal derivatives were also modeled. Trials on which participants failed to respond and time-points with excessive motion (>1.5mm) were modeled with two separate regressors and dropped from analyses.

Within each participant, each run was modeled separately and then combined using a fixed-effects model. Data across participants (within-group and between-group) were analyzed using a random-effects model. We used Monte Carlo simulations as implemented in FSL to correct for multiple comparisons across the whole cortex. For within-group results, on each permutation iteration, voxel values signs across the brain are flipped (e.g. 4.5 to −4.5) for a random subset of subjects and the subsequent group map is thresholded at a cluster-forming threshold (p<0.01) (Winkler et al. 2014). The size of largest number of contiguous vertices is then entered into a null distribution and clusters from our true results that lie in the top 5% (alpha of p<0.05) of this distribution pass the cluster-correction. The correction procedure for between-group results was similar except group labels were permuted rather than voxel value signs (Winkler et al. 2014).

Math-responsive regions of interest (ROIs) in the intraparietal sulcus (IPS) were defined within an anatomical IPS search-space, using a leave-one-run-out procedure. Using all but one run, ROIs were defined by taking the top 20 vertices within the search-space with the greatest math>language effect (Destrieux et al. 2010). Percent signal change (PSC) for all four math conditions and the language condition was then extracted from the left out run using finite impulse response modeling (Lindquist et al. 2009). This procedure was repeated iteratively until PSC was extracted from every run and the results were averaged across the iterations.

We then looked for an effect of digit-number and algebraic complexity, which are orthogonal to the math>sentence contrast used for ROI definition. We also tested selectivity for math over language by comparing the math and sentence conditions (note that independent data were used to define math>sentence ROIs). Under the null hypothesis, the vertices that show the math>sentence effect in the runs used to define the ROI are random, and would not be expected to show the effect in held out run.

Within visual cortex, we looked at activity in math-responsive rMOG, which has previously been observed to respond to numerical information in congenitally blind individuals (Kanjlia et al. 2016). Math-responsive ROIs in the visual cortex were defined as follows: for each congenitally blind and sighted subject, a search-space was created by taking the rMOG cluster that responded to the math>language contrast in CB>S (p<0.0001, uncorrected). Each congenitally blind and sighted participant did not contribute to the creation of his or her own search-space. Each congenitally blind and sighted participant was “left out,” iteratively, and his or her search-space was created based on functional data from the remaining subjects. Since search-space definition procedure was independent of the adult-onset blind group, the same search-space was used for all adult-onset blind subjects (all CB>S, math>language, p<0.0001, uncorrected). Functional ROIs were then defined within the search-space in every subject using the leave-one-run-out procedure described above. Additionally, we looked at responses in V1 because this is the first cortical stage of visual processing. The functional reorganization of this region is of particular interest and has been investigated in many prior studies of sensitive periods in visual cortex plasticity (Cohen et al. 1999; Bedny et al. 2012; Collignon et al. 2013b).

All analyses with multiple measures per subject treated subjects as a random-effect. Paired t-tests were used to compare means within a group and unpaired t-tests were used when comparing means across groups. All t-tests were two-tailed.

Correlations with duration of blindness were conducted including only adult-onset blind participants who lost their vision abruptly (within 2 years, n=7; see Table 1) because blindness duration is less clearly defined when vision is lost progressively.

### Resting-state functional connectivity analysis

Resting-state correlations will be referred to as functional connectivity henceforth. Resting-state data were analyzed using CONN v.17 Functional Connectivity Toolbox (Whitfield-Gabrieli and Nieto-Castanon 2012). Functional data were linearly detrended by including a linear regressor in the general linear model to remove low-frequency drift. Data were despiked by applying a hyperbolic tangent “squashing” function to data from every time point. Data were band-pass filtered (0.008-0.1 Hz) and signal from white mater and cerebrospinal fluid were regressed out. Functional data were smoothed 23 diffusion steps (corresponding to ∼6mm smoothing in volume) (Hagler et al. 2006). Fisher-transformed *r* values were used for statistical analyses.

ROI-to-ROI resting-state functional connectivity analyses were conducted in the right hemisphere, since task-based effects were right-lateralized. Search-spaces were defined across groups and group-specific (congenitally blind, adult-onset blind and sighted) ROIs were defined within these search-spaces. To avoid biasing search-space definition to groups with a larger sample size, we used data from all 13 adult-onset blind participants, the first 13 congenitally blind and first 13 sighted participants to define search-spaces. This subsample of 39 participants was entered into a single random-effects model to find prefrontal math (math>language) and language (language>math) responsive areas common across groups (p<0.01, uncorrected). Within these broad regions, math- and language-responsive prefrontal ROI’s were defined separately for each group (using all participants for that group) by taking the top 250 vertices with the greatest response to the math>language and language>math contrast, respectively. Math-responsive IPS ROI’s were defined for each group by taking the top 250 vertices with the greatest math>language effect within anatomically defined IPS search-space (Destrieux et al. 2010).

Math- and language-responsive ROIs in the visual cortex could only be defined in the congenitally blind group and thus CB ROIs were used for all groups. A cluster in dorsal occipital cortex that responded to the math>language contrast in CB>S served as the math-responsive visual cortex ROI (p<0.01, uncorrected). A cluster in ventral occipito-temporal cortex (within occipital lobe mask) that responded to the language>math contrast in CB>S served as the language-responsive visual cortex ROI (p<0.01, uncorrected).

## Results

### Behavioral Results

In adult onset blind participants, accuracy and response times were similar across math and sentence conditions (accuracy: t(12)=0.58, p=0.57; response times: t(12)=1.02, p=0.33) (Fig. 1). As previously reported for congenitally blind and sighted individuals (Kanjlia et al. 2016), adult-onset blind individuals were faster and more accurate on trials with single-digit than double-digit math equations (digit-number by algebraic complexity repeated measures ANOVA; main effect of digit-number on accuracy: F(1,12)=9.88, p=0.008; main effect of digit-number on response times: F(1,12)=9.00, p=0.01) (Fig. 1). Similarly, adult-onset blind individuals were faster more accurate on trials with algebraically simple math problems than algebraically complex problems (main effect of algebraic complexity on accuracy: F(1,12)=21.41,p=0.001; main effect of algebraic complexity on response times: F(1,12)= 15.82, p=0.002).

The adult-onset blind group was less accurate than the congenitally blind and sighted group across the math and language tasks (task by group repeated measures ANOVA: main effect of group (AB vs CB): F(1,31)=6.96, p=0.01; main effect of group (AB vs. S): F(1,30)=5.37, p=0.03). The adult-onset blind group was slightly less accurate than the sighted group on math trials (t(30)=2.1, p=0.04) and less accurate on sentence trials relative to both of the other groups (AB vs. CB: t(31)=3.60, p=0.001; AB vs. S: t(30)=2.03, p=0.051). Adult-onset blind individuals were marginally slower to respond on sentence trials compared to the congenitally blind group (AB vs. CB: t(31)=-2.00, p=0.06) and slower on math trials compared to the sighted group (AB vs. S: t(30)=-2.30, p=0.03). All other comparisons were not significant (p>0.05; Supplementary Table 1).

### Similar fronto-parietal responses in adult-onset blind, congenitally blind and sighted groups

All three groups showed similar responses in fronto-parietal cortices for the math>language contrast (p<0.05, cluster-corrected, Fig. 2, Fig. 3, Supplementary Table 2). ROI analyses show that, like the IPS of congenitally blind and sighted individuals, the IPS of adult-onset blind individuals responded more to the math than the language task (AB group, hemisphere by task repeated-measures ANOVA; main effect of task (math vs. language): F(1,12)=187.91, p<0.001; hemisphere by task interaction: F(1,12)=14.71, p=0.002; Supplementary Table 3) and showed the same sensitivity to digit-number (hemisphere by digit-number by algebraic complexity repeated-measures ANOVA; main effect of digit-number in AB group: F(1,12)=14.38, p=0.003; digit-number by group (AB vs. S) interaction: F(1,30)=0.95, p=0.34; digit-number by group (AB vs. CB) interaction: F(1,31)=0.002, p=0.96; Supplementary Table 3). The adult-onset blind group did not show an effect of algebraic complexity (AB group: F(1,12)=0.20, p=0.66) in the IPS. The effect of algebraic complexity was not different across adult-onset blind and congenitally blind groups but was slightly larger in the sighted group compared to the adult-onset blind group (algebraic complexity by group (AB vs. CB) interaction: F(1,31)=0.84, p=0.37; algebraic complexity by group (AB vs. S) interaction: F(1,30)=3.18, p=0.09).

**Figure 2.**
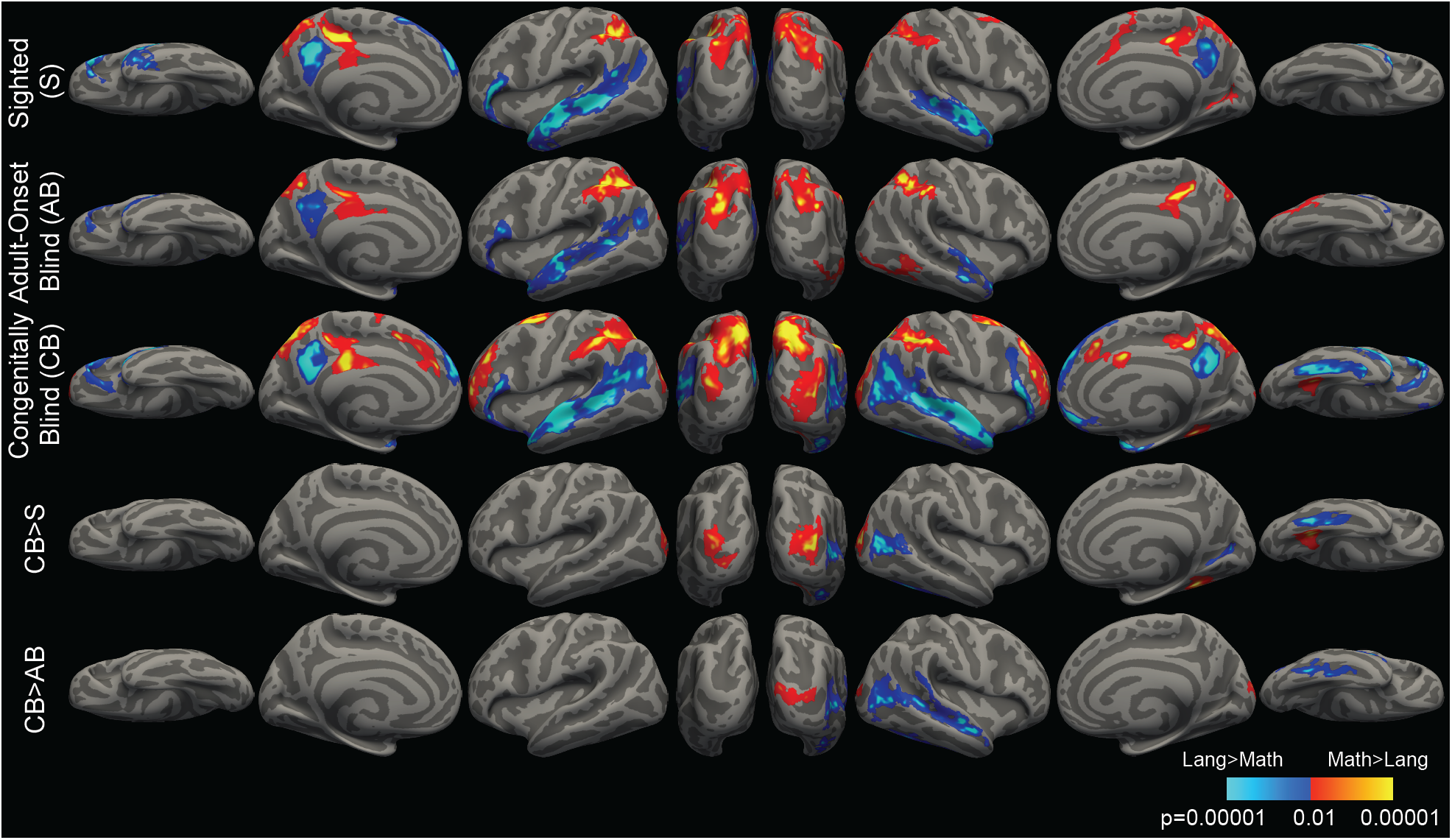
Whole-Cortex Responses to Math and Language. Brain regions active for math > language (warm colors) and language > math (cool colors) (p < 0.05, cluster corrected).

**Figure 3.**
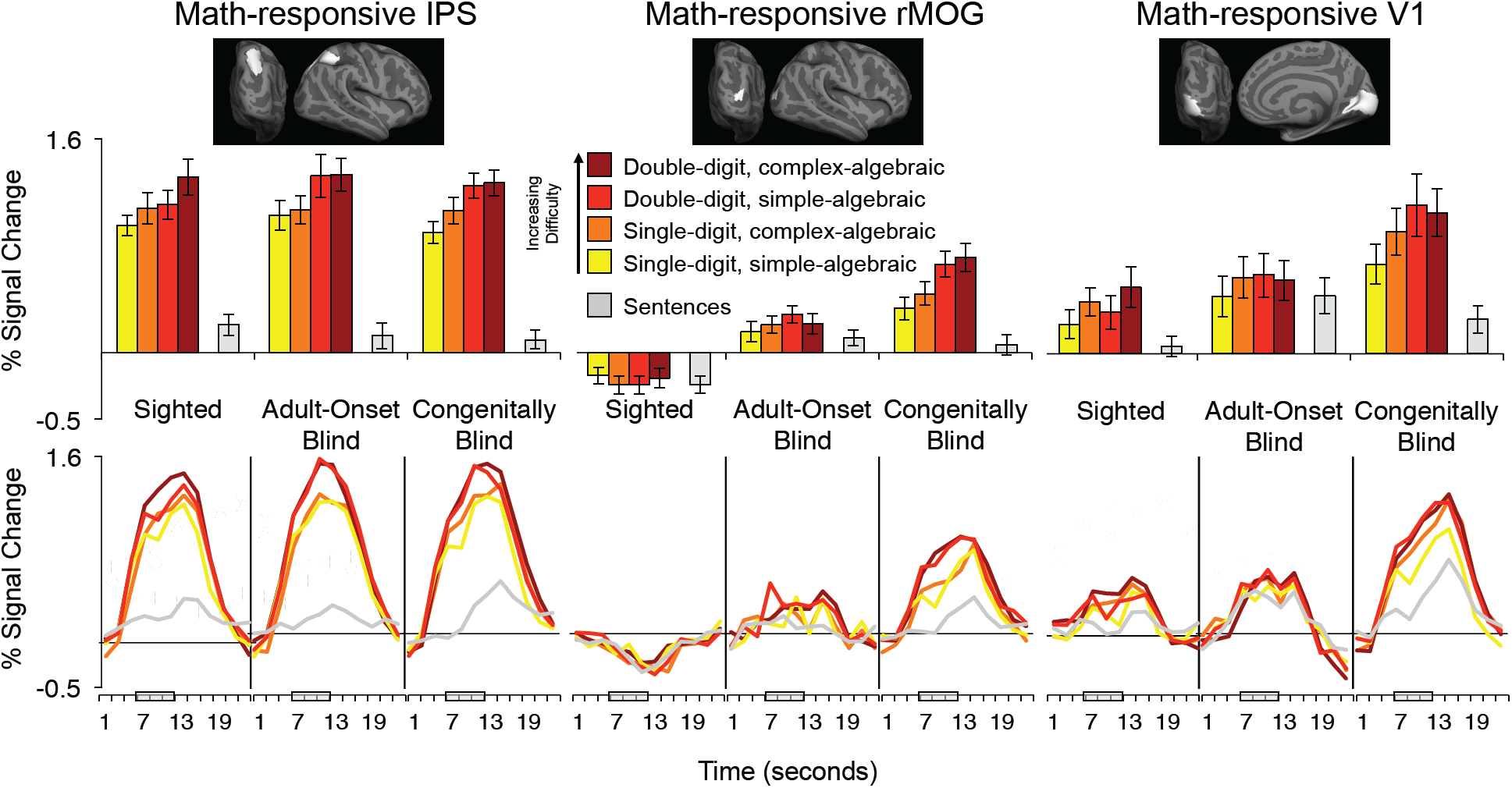
Math and Language Activity in IPS, rMOG and V1. Responses to math equations by difficulty in math-responsive IPS (left), math-responsive rMOG (middle) and math-responsive V1 (right). Percent signal change relative to rest was extracted from individual-subject ROIs defined within IPS, rMOG and V1 search-spaces. Adult-onset blind search-spaces displayed at the top. IPS and V1 results are averaged across left and right hemispheres. Error bars represent standard error of the mean.

### Different visual cortex sensitivity to higher-cognitive functions in congenitally blind as opposed to adult-onset blind and sighted groups

Relative to the sighted, congenitally blind but not adult-onset blind participants activated several regions within “visual” cortex during math calculation versus sentence comprehension and vice versa: in whole-cortex analyses, the rMOG was more active for math than language while the rVOT and right lateral occipital cortex (rLO) were more active for language than math (Fig. 2). Although some visual cortex activity was observed in the within-group analysis of the adult-onset blind group, this activity was focused around the location of the so-called visual number-form area (VNFA), which has previously been shown to respond to numerical tasks in sighted individuals and was also observed in the sighted group at a reduced statistical threshold in the present study (Abboud et al. 2015). Direct comparison of congenitally blind and adult-onset blind participants revealed greater rMOG activity in the congenitally blind for the math>language contrast and greater right rLO activity in the congenitally blind for language>math contrast (Fig. 2, CB>AB, math>language, p<0.05, cluster-corrected).

In ROI analyses, overall response to all math and language conditions in rMOG was greater in both congenitally and adult-onset blind groups compared to the sighted group (CB vs. S: t(37)=6.30, p<0.001; AB vs. S: t(30)=4.73, p<0.001; Fig. 3). rMOG response to all stimuli was marginally higher in the congenitally blind group than the adult-onset blind group (t(31)=1.94, p=0.06). Selectivity for mathematical stimuli over sentence stimuli was also significantly larger in congenitally blind as compared to the adult-onset blind group (CB vs. AB; task by group interaction: F(1,31)=10.72, p=0.003). However, the rMOG showed a larger response to mathematical stimuli over sentence stimuli in adult-onset blind individuals as well (math vs. language, AB: t(12)=2.28, p=0.04; CB: t(19)=5.5, p<0.001). There was no difference in rMOG selectivity for math over language stimuli across adult-onset blind and sighted individuals (AB vs. S; task by group interaction: F(1,30)=1.27, p=0.27).

Similarly, the effect of digit-number was larger in the congenitally blind than the adult-onset blind group (digit-number by group interaction: F(1,31)=9.58, p=0.004). There was a marginal difference in the algebraic complexity effect across congenitally blind and adult-onset blind groups (algebraic complexity by group interaction: F(1,31)=3.28, p=0.08). The rMOG of the adult-onset blind was not different from that of the sighted in its sensitivity to either math difficulty manipulation (digit-number by group interaction: F(1,30)=2.88, p=0.10; algebraic complexity by group interaction: F(1,30)=0.004, p=0.95). Within the adult-onset blind group, the rMOG did not show sensitivity to either digit-number or algebraic complexity (AB group, digit-number by algebraic complexity ANOVA; main effect of digit-number: F(1,12)=2.90, p=0.12; main effect of algebraic complexity: F(1,12)=0.06, p=0.82; Supplementary Table 3).

In V1, selectivity for mathematical stimuli over sentence stimuli was stronger in the congenitally blind than the adult-onset blind group and marginally larger in the sighted than the adult-onset blind group (hemisphere by task by group repeated measures ANOVA: CB vs. AB: F(1,31)=18.87, p<0.001; AB vs. S: F(1,30)=3.43, p=0.07; Fig. 3; Supplementary Table 3). The effect of digit-number was larger in the congenitally blind than the adult-onset blind group (hemisphere by digit-number by algebraic complexity by group repeated measures ANOVA: digit-number by group (CB vs. AB) interaction: F(1,31)=4.18, p=0.05). Interestingly, the sighted group showed a significant effect of algebraic complexity in V1 (main effect of algebraic complexity: F(1,18)=10.67, p=0.004; main effect of digit-number: F(1,18)=1.70, p=0.21). By contrast, adult-onset blind individuals show no sensitivity to digit-number or algebraic complexity (main effect of digit-number: F(1,12)=1.16, p=0.30; main effect of algebraic complexity: F(1,12)=0.90, p=0.36; S vs. AB algebraic complexity by group interaction: F(1,30)=2.58, p=0.12).

Notably, selectivity for math (% signal change for mathematical stimuli-language stimuli) in the rMOG and V1 was not predicted by duration of blindness among adult-onset blind participants with abrupt vision loss (see Methods) or congenitally blind participants (i.e. age) (AB rMOG: R2=0.02, p=0.79; AB V1: R2=0.17, p=0.36; CB rMOG: R2=0.05, p=0.34; CB V1: R2=0.00, p=0.91). Similarly, there was no correlation between blindness duration and the size of the math difficulty effect (% signal change for hardest math condition – easiest math condition) in either the rMOG or V1 of the AB or CB (AB rMOG: R2=0.46, p=0.09; AB V1: R2=0.08, p=0.53; CB rMOG: R2=0.01, p=0.76; CB V1: R2=0.03, p=0.45).

### Functional connectivity between “visual” cortices and fronto-parietal cortices in adult-onset blindness

In congenital blindness, visual cortices become more correlated at rest with parietal and prefrontal cortices (Kanjlia et al. 2016). We confirm this effect with larger sample of congenitally blind participants: math-responsive rMOG and language-responsive rVOT were more correlated with the IPS, rDLPFC and rIFG in the congenitally blind as opposed to sighted (main effect of group (CB vs. S) connectivity of visual cortex to IPS: F(1,65)=24.49, p<0.001; main effect of group connectivity of visual cortex to prefrontal cortices (rDLPFC and rIFG): F(1,65)=16.11, p<0.001; Fig. 4).

**Figure 4.**
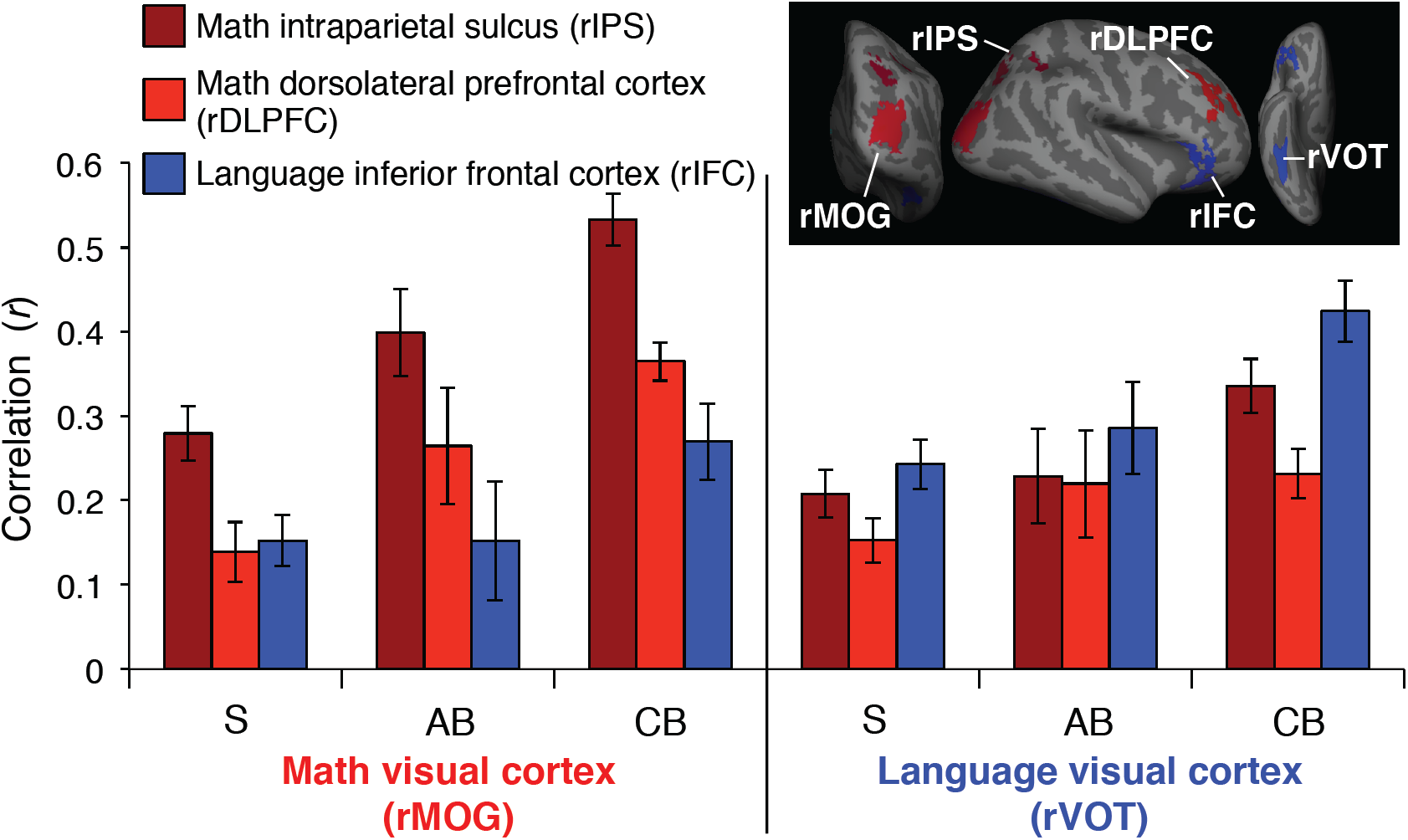
Resting-State Functional Connectivity Between Occipital and Fronto-Parietal Networks. Resting-state correlations between math-responsive (left) and language-responsive (right) visual cortices and fronto-parietal math network (red) and inferior frontal language region (blue). ROIs for sighted group shown above (see Supplementary Fig. 1 for congenitally blind and adult-onset blind group ROIs). Error bars show standard error of the mean.

As previously reported, we found that increases in functional connectivity among congenitally blind individuals are network-specific. Math-responsive rMOG but not language responsive rVOT shows elevated resting-state correlations with math-responsive rIPS (Fig. 4; seed (rMOG vs. rVOT) by group (CB vs. S) interaction: F(1,65)=5.32, p=0.02). Similarly, while math-responsive visual cortex (rMOG) becomes more correlated with math-responsive portions of prefrontal cortex (rDLPFC), language-responsive visual cortex (VOT) becomes more correlated with inferior frontal language areas (seed (rMOG vs. rVOT) by ROI (rDLFPC vs. rIFC) by group (CB vs. S) interaction: F(1,65)=12.39, p=0.001).

Although the specialization of functional connectivity is stronger in the congenitally blind group, within-group analyses showed that both for the congenitally blind and for the sighted, within-network correlations (math visual cortex to math prefrontal cortex) are higher than between network correlations (math visual cortex to language prefrontal cortex) (seed by ROI interaction in CB group: F(1,23)=23.41, p<0.001; and sighted group: F(1,42)=6.57, p=0.01). This effect of resting-state functional connectivity specialization among the sighted has not previously been observed, likely due to smaller samples of blindfolded sighted participants in previous studies (Kanjlia et al. 2016).

Among the adult-onset blind group, resting-state functional connectivity of visual cortices show an intermediate pattern between the sighted and congenitally blind groups (Fig. 4).

Overall magnitude of correlation between visual cortices and the IPS and visual cortices and prefrontal cortices is marginally lower in the adult-onset blind group, compared to the congenitally blind and is not different from the sighted (connectivity with IPS, seed (rMOG vs. rVOT) by group (AB vs. CB) repeated measures ANOVA, main effect of group: F(1,34)=6.14, p=0.02; connectivity with prefrontal cortices, seed (rMOG vs. rVOT) by ROI (rDLPFC vs. rIFC) by group (AB vs. CB) repeated measures ANOVA, main effect of group: F(1,34)=3.25, p=0.08; connectivity with IPS, seed by group (AB vs. S) ANOVA, main effect of group: F(1,53)=1.68, p=0.20; connectivity with prefrontal cortices, seed by ROI by group (AB vs. S); main effect of group: F(1,53)=1.15, p=0.29).

Resting-state correlations of visual cortices among the adult-onset blind group show clear network selectivity: activity of math-responsive visual cortex (rMOG) is more correlated with math-responsive parietal (rIPS) and prefrontal (rDLPFC), whereas activity of language-responsive visual cortex (rVOT) is more correlated with language-responsive inferior frontal cortex (rIFC) (within adult-onset blind group; connectivity with IPS, effect of seed (rMOG vs. rVOT): t(11)=3.52, p=0.005; connectivity with prefrontal cortices, seed (rMOG vs. rVOT) by ROI (rDLPFC vs. rIFC) interaction: F(1,11)=7.81, p=0.02).

Selectivity of functional connectivity across number and language networks in adult-onset blindness did not differ from either the congenitally-blind or sighted groups (connectivity with IPS, seed (rMOG vs. rVOT) by group (AB vs. CB) interaction: F(1,34)=0.17, p=0.68; connectivity with prefrontal cortices, seed by ROI (rDLPFC vs. rIFC) by group (AB vs. CB) interaction: F(1,34)=1.28, p=0.27; connectivity with IPS, seed by group (AB vs. S) interaction: F(1,53)=2.00, p=0.16; connectivity with prefrontal cortices, seed by ROI by group (AB vs. S) interaction: F(1,53)=2.40, p=0.13).

Notably, among adult-onset blind individuals with abrupt vision loss (see Methods), resting-state functional connectivity between rMOG and rIPS but not rPFC was significantly correlated with blindness duration since reaching one’s current level of vision (rIPS: R2=0.72, p=0.02; rPFC: R2=0.14, p=0.42).

## Discussion

### Sensitive period for cognitive repurposing in visual cortex

We find that the capacity of cortex to take on novel cognitive functions narrows after development. In congenital blindness, different visual cortex regions become specialized for numerical as opposed to linguistic processing and BOLD signal in these regions increases with cognitive load (Bedny et al. 2011; Lane et al. 2015; Kanjlia et al. 2016). A dorsal occipital area (rMOG) and parts of V1 are more responsive to math equations than sentences and activity increases with the difficulty of math equations in congenitally blind but not sighted participants (Kanjlia et al. 2016). By contrast, regions in ventral occipito-temporal cortex (VOT) and lateral occipital cortex (LOC) are more responsive to sentences (Bedny et al. 2011; Lane et al. 2015; Kim et al. 2017).

Here we report that this type of cognitive repurposing is qualitatively different in individuals who lose their vision as adults. In adult-onset blindness (blind at age 17 or later), there is less regional specialization within visual cortex (i.e. for numerical and linguistic processing). Instead, the “visual” cortex shows an above rest response across cognitive tasks and conditions. Crucially, relative to the congenitally blind, visual cortices of adult-onset blind participants show less sensitivity to mathematical difficulty (i.e. cognitive load). This is despite the fact that, in adult-onset and congenitally blind participants alike, the overall amount of visual cortex activity during auditory tasks is elevated relative to rest, as are resting-state correlations of visual cortex with fronto-parietal networks (Bedny et al. 2012; Collignon et al. 2013b).

Differences in the functional profile of visual cortex cross the adult-onset and congenitally blind groups do not appear to be related to the blindness duration, since neither the selectivity of the visual cortex for math equations nor its response to equation-difficulty increased with blindness duration among the adult-onset or congenitally blind participants. As with any null result it remains possible that an effect of blindness duration does exist in the population and was not detected in the current study, perhaps due to insufficient power. However, the present results suggest that any putative effect of blindness duration coexists with a more robust effect of age of blindness onset.

Why might the recruitment of visual cortex for higher-cognitive functions be limited to a sensitive period during development? One possibility is that cognitive specialization of cortex requires circuit-internal structural changes that are uniquely possible during sensitive periods in development. As noted in the introduction, studies in animals suggest that dendritic spine formation, spine elimination and axon retraction are enhanced during sensitive periods (Hensch, 2004; Hensch, 2005; Hensch, 2005; Maurer and Hensch, 2012). Sensitive period closure coincides with formation of molecular “brakes,” such as perineuronal nets, which dampen plasticity (Pizzorusso 2002; Bavelier et al. 2010). Enhanced levels of structural flexibility in visual cortex during sensitive periods may enable it to acquire non-visual cognitive functions in those who are blind from birth and early blind. According to this hypothesis, cognitive repurposing of visual cortex depends on sensitive period neurophysiology, which declines over the first few years of life in humans (Maurer and Hensch 2012). Alternatively, establishing one set of representations (e.g. visual) could block cortex from representing other content (e.g. number). If so, repurposing of visual cortex is only possible in individuals who are “visually naïve.”

In support of the structural flexibility hypothesis, previous studies provide some evidence for gradual decline in cross-modal responses with age of blindness onset. For example, the amount of visual cortex activity in early blind individuals during Braille and spoken language tasks is intermediate between that of congenitally and adult-onset blind individuals (Cohen et al. 1999; Sadato et al. 2002; Burton et al. 2003). However, these studies compare non-visual tasks to rest and the current data suggest that responses to higher-cognitive information in visual cortex have a different developmental time-course than responses to non-visual stimulation in general. Future work should test the generalizability of the present findings to tactile tasks, such as Braille reading, and ask whether the capacity of visual cortex to specialize for *specific* cognitive operations declines gradually over childhood or abruptly after birth.

A further question raised by the current findings concerns the cognitive and behavioral significance of visual cortex activity in adult-onset blindness. As noted in the introduction, sensory cortices can assume new, behaviorally relevant functions even in adulthood. Amputation of a limb causes deaffrented somatosensory cortices to respond to body parts represented by neighboring regions and there is some evidence that these responses are behaviorally relevant (Pascual-Leone et al. 1996; Röricht et al. 1999). However, in such cases, functional repurposing occurs within a modality (Masuda et al. 2008, 2010; Baseler et al. 2011; Srihasam et al. 2012; Lemos et al. 2016). Whether adult cortex can repurpose across modalities remains an open question. In the current study, visual cortex activity during auditory tasks may not be cognitively or behaviorally relevant in adult-onset blindness. Consistent with this possibility, even though visual cortices of congenitally and adult-onset blind individuals are active during Braille reading tasks, TMS to the visual cortex impairs Braille reading only in those who are congenitally blind (Cohen et al. 1999). Alternatively, the visual cortex of adult-onset blind individuals may take on non-visual cognitive functions that are different from those it takes on in congenital blindness, perhaps functions that are easier for mature cortex to acquire. Under this view, adult cortex can repurpose but only within a narrow cognitive range.

It is worth noting that although cognitive repurposing of visual cortex in the adult onset blind group is greatly reduced relative to congenitally blind individuals, the visual cortex nevertheless does change its function to some degree even in adult-onset blindness relative to the sighted. In the rMOG there was a small but significant preference for math over language stimuli in the adult-onset blind group but not in the sighted group. This effect was weaker than what was found in the congenitally blind group and, unlike in the congenitally blind group, there was no effect of cognitive load. In V1, there was a small but significant difference between math and language in the sighted group that was actually absent in the adult-onset blind group. This finding is consistent with some previously observed non-visual responses in the V1 of the sighted and could indicate the loss of this response in adult-onset blindness (Merabet et al. 2006; Sathian and Stilla 2011; Vetter et al. 2014). Together these results suggest that blindness in adulthood does, in fact, change the function of the visual cortex, but not in the same way or to the same degree as blindness at birth. We hypothesize that there is a sensitive period for cortex to assume a specific new cognitive function, but no sensitive period for functional change per se. Future work is needed to understand the capacity of the “visual” cortex to repurpose for cognitive functions other than those studied here and to determine how sensitive periods vary by visual region.

Exactly what defines the cognitive potential of cortex in adulthood and what distinguishes it from the cognitive range of developing cortex remains an open question for future research. Notably, even though the present findings suggest that the cognitive range of adult cortex is naturally restricted, pharmacological and even targeted behavioral interventions (e.g. sensory deprivation or environmental enrichment), can “reopen” sensitive periods (Putignano et al. 2007; Baroncelli et al. 2010; Bavelier et al. 2010; Maya Vetencourt et al. 2011; Spolidoro et al. 2011). Therefore the existence of such windows of sensitivity is better viewed as a time of greatest neurocognitive flexibly, rather than as a unique and immutable window for change.

### Functional connectivity of visual cortices changes, even in adult-onset blindness

Although we find that the visual cortices of adult-onset blind individuals do not take on the same cognitive functions as those of congenitally blind individuals, blindness in adulthood still changes the functional properties of visual cortex: resting-state correlations between visual cortices and the fronto-parietal number network increase.

These findings are consistent with a recent study that found increased resting-state correlations between visual cortices and Broca’s area in individuals who became totally blind after the age of 21 due to retinitis pigmentosa compared to sighted individuals (Sabbah et al. 2016). Interestingly, the same study found a similar increase in functional fronto-occipital connectivity even in the case of partial vision loss (Sabbah et al. 2016). Together these findings suggest that functional connectivity of visual cortex remains modifiable into adulthood. It is worth noting, however, that we and others have found that resting-state correlations between visual cortex and higher-cognitive networks are lower in those who are adult-onset as compared to congenitally blind (Bedny et al. 2010; Butt et al. 2013). In this respect the adult-onset blind group is intermediate between what is observed in congenital blindness and in the blindfolded sighted group. Therefore, the flexibility of the adult brain, even in the case of functional connectivity, is not quite as extensive as that of the juvenile brain.

Importantly, in adult-onset blind individuals, visual cortices not only demonstrate increased resting-state correlations with fronto-parietal networks overall, but exhibit region-specific increases with different fronto-parietal functional networks, similar to what is found in congenital blindness (Kanjlia et al. 2016). In particular, visual areas that respond to math equations in the congenitally blind group are correlated with the fronto-parietal number network in the adult-onset blind group. By contrast, those that respond to language in congenital blindness are correlated with inferior frontal language areas in the adult-onset blind group. This pattern is surprising, given that adult-onset blind individuals do not show sub-specialization of the visual cortex for math and language processing in task-based data. A small but significant functional connectivity dissociation among visual areas was observed even in blindfolded sighted controls.

A key open question concerns how resting-state correlations and task-based functional selectivity relates to the underlying anatomical connectivity patterns of visual cortex. One possibility is that anatomical connectivity biases across visual cortex networks give rise to both the resting-state and the task-based selectivity patterns. According to this idea, in sighted and blind infants alike, there is stronger anatomical connectivity between the rMOG region of visual cortex and the fronto-parietal number network on the one hand, and the rVOT region of the visual cortex and the fronto-temporal language network on the other. In the sighted, this anatomical pattern gives rise to some region-specific fronto-occipital synchrony but does not lead to the specialization for number and language in the visual cortex, because non-visual inputs are dwarfed by bottom-up inputs from the visual pathway. By contrast, in congenital blindness, this anatomical bias leads to both functional synchronization at rest and sensitivity to language and number in different “visual” areas. Finally, adult-onset blindness leads only to the up-regulation of resting-state correlations between these anatomically connected regions but not to task-based responses to language and number.

At present the above hypothesis is speculative and remains to be tested. Previous studies have shown that functional connectivity reflects a complex combination of anatomical and functional factors (Damoiseaux and Greicius 2009; Greicius et al. 2009; Honey et al. 2009). Cortical regions that have strong long-range anatomical connections tend to have stronger functional connectivity, however, regions can be synchronized through intermediary areas and need not have direct anatomical connections (Damoiseaux and Greicius 2009; Greicius et al. 2009; Honey et al. 2009). Furthermore, resting-state connectivity partly reflects a past history of co-activation above and beyond the strength of anatomical connectivity (Lewis et al. 2009). Therefore, a pair of regions with similar amounts of long-range anatomical connectivity can exhibit different resting-state correlation patterns across populations with different life histories. This point is illustrated in studies of blindness that find enhanced fronto-occipital synchrony in blind relative to sighted individuals despite no clear increases in anatomical connectivity (Shimony et al. 2006; Liu et al. 2007; Yu et al. 2008). In addition to influencing resting-state functional connectivity patterns, anatomical biases have also been shown to determine the localization of cognitive functions. For example, in young children, the visual word form area (VWFA) has strong anatomical connectivity with fronto-temporal language networks even before literacy (Dehaene et al. 2015). Furthermore, the location of these anatomical connections within the ventral occipito-temporal cortex predicts individual differences in the future location of letter and word responses in the ventral stream (Saygin et al. 2016). Notably, in congenital blindness, the VWFA is one of the “visual” areas that becomes responsive to high level linguistic content (i.e. grammar) (Lane et al. 2015; Kim et al. 2017). Such evidence provides general support for the idea that anatomical connectivity predicts functional synchrony and task-based responses. Whether it does so in the specialization of visual cortex for number as opposed to language remains to be tested. Future work could use diffusion tractography imaging (DTI) to directly compare structural connectivity of math- and language-responsive visual areas between sighted and blind individuals (Pascual-Leone et al. 2005; Wang et al. 2015).

The present results are consistent with prior evidence that resting-state connectivity patterns relate to functional specialization of cortex. However, the current findings also highlight a dissociation between long-range resting-state connectivity and local functional properties. The “visual” cortex of adult-onset and congenitally blind participants show similar resting-state functional connectivity yet different task-based responses. One interpretation of these results is that communication between cortical areas is necessary but not sufficient for functional specialization. If inputs reach a cortical area only after a sensitive period has closed, cognitive specialization of that local circuit may fail to occur despite receiving relevant information.

### Conclusions

In summary, we find that blindness at any age causes visual cortices to become synchronized with multiple different higher-cognitive fronto-parietal networks in a region-specific manner. However, the visual cortices of adult-onset and congenitally blind adults show different capacity to take on higher-cognitive functions. These findings suggest that the capacity of cortex to take on novel functions is restricted to sensitive periods of development, possibly due to local cortical neurophysiology.

## Acknowledgements

We thank the F. M. Kirby Research Center for Functional Brain Imaging at the Kennedy Krieger Institute for their assistance with data collection; Connor Lane for assisting with data collection; and the Baltimore and Washington, DC, blind communities. Research reported in this publication was supported by the National Eye Institute of the National Institutes of Health under award number R01EY027352 and the Science of Learning Institute at Johns Hopkins University under award #80034917. The content is solely the responsibility of the authors and does not necessarily represent the official views of the National Institutes of Health.

## Supplementary Material

**Supplementary Table 1.** Behavioral Results.

|  | Accuracy |  | Response Time |  |
| --- | --- | --- | --- | --- |
|  | AB vs. CB | AB vs. S | AB vs. CB | AB vs. S |
| Effect of Digit-Number | F(1,31)=0.002, p=0.96 | F(1,30)=2.37, p=0.13 | F(1,31)=1.35, p=0.25 | F(1,30)=0.66, p=0.42 |
| Effect of Algebraic Complexity | F(1,31)=3.01, p=0.09 | F(1,30)=6.51, p=0.02 | F(1,31)=0.28, p=0.60 | F(1,30)=5.12, p=0.03 |
| Effect of Task | F(1,31)=2.57, p=0.12 | F(1,30)=0.31, p=0.59 | F(1,31)=1.27, p=0.27 | F(1,30)=1.46, p=0.24 |
| Math | t(31)=1.41, p=0.17 | t(30)=2.1, p=0.04 | t(31)=-1.85, p=0.07 | t(30)=1.67, p=0.11 |
| Sentences | t(31)=3.60, p=0.001 | t(30)=2.03, p=0.051 | t(31)=-2.54, p=0.02 | t(30)=0.87, p=0.39 |

**Supplementary Table 2.** Brain regions active during math and language tasks.

| Brain regions active for math > language | x | y | z | Peak t | mm2 | Pcluster |
| --- | --- | --- | --- | --- | --- | --- |
| **Adult-Onset Blind Group** |  |  |  |  |  |  |
| Left postcentral sulcus | 40 | -40 | 37 | 13.56 | 3247.83 | 0.0008 |
| Left intraparietal sulcus and transverse parietal sulci | 31 | -65 | 34 | 10.53 |  |  |
| Left precuneus | 7 | -65 | 50 | 9.05 |  |  |
| Left marginal branch of the cingulate sulcus | 7 | -33 | 43 | 6.84 | 671.49 | 0.043 |
| Right intraparietal sulcus and transverse parietal sulci | 29 | -51 | 44 | 12.67 | 2274.23 | 0.0002 |
| Right supramarginal gyrus | 56 | -41 | 42 | 10.78 |  |  |
| Right middle occipital gyrus | 35 | -79 | 34 | 9.01 |  |  |
| Right superior occipital sulcus and transverse occipital sulcus | 28 | -64 | 29 | 7.5 |  |  |
| Right superior parietal lobule | 17 | -68 | 54 | 7.09 |  |  |
| Right inferior temporal sulcus | 55 | -53 | -4 | 5.98 | 590.13 | 0.033 |
| Right inferior occipital gyrus and sulcus | 45 | -82 | -9 | 5.08 |  |  |
| Right marginal branch of the cingulate sulcus | 7 | -38 | 43 | 11.66 | 579.53 | 0.0332 |
| **Congenitally Blind Group** |  |  |  |  |  |  |
| Left superior parietal lobule | -17 | -70 | 45 | 9.62 | 4297.49 | 0.0002 |
| Left supramarginal gyrus | -52 | -39 | 47 | 7.96 |  |  |
| Left middle occipital gyrus | -38 | -88 | 16 | 6.25 |  |  |
| Left middle frontal gyrus | -39 | 50 | 9 | 8.39 | 1703.1 | 0.003 |
| Left middle frontal gyrus | -44 | 31 | 30 | 6.46 |  |  |
| Left fronto-marginal gyrus and sulcus | -23 | 56 | -7 | 5.02 |  |  |
| Left superior frontal sulcus | -21 | 7 | 50 | 9.51 | 1127.99 | 0.0086 |
| Left posterior-dorsal part of the cingulate gyrus | -6 | -30 | 29 | 6.88 | 871.01 | 0.016 |
| Left marginal branch of the cingulate sulcus | -11 | -41 | 45 | 6.46 |  |  |
| Left middle-anterior part of the cingulate gyrus and sulcus | -8 | 8 | 45 | 6.4 | 642.01 | 0.0286 |
| Left anterior part of the cingulate gyrus and sulcus | -9 | 35 | 26 | 6.32 |  |  |
| Right sulcus intermedius primus | 43 | -44 | 36 | 10.3 | 3551.84 | 0.002 |
| Right intraparietal sulcus and transverse parietal sulci | 19 | -63 | 53 | 9.07 |  |  |
| Right marginal branch of the cingulate sulcus | 7 | -41 | 44 | 7.95 |  |  |
| Right middle frontal gyrus | 38 | 27 | 39 | 7.36 | 1591.46 | 0.0068 |
| Right inferior frontal sulcus | 43 | 33 | 20 | 7.12 |  |  |
| Right middle frontal sulcus | 30 | 50 | 0 | 5.89 |  |  |
| Right middle occipital sulcus and lunatus sulcus | 33 | -82 | 9 | 6.83 | 1204.35 | 0.0092 |
| Right superior frontal sulcus | 28 | 6 | 51 | 8.29 | 925.6 | 0.0138 |
| Right middle-posterior part of the cingulate gyrus and sulcus | 3 | 3 | 34 | 7.45 | 570.11 | 0.0342 |
| Right superior frontal gyrus | 6 | 23 | 43 | 6.62 |  |  |
| Right medial occipito-temporal sulcus and lingual sulcus | 31 | -45 | -14 | 6.04 | 455.74 | 0.0476 |
| **Sighted Group** |  |  |  |  |  |  |
| Left intraparietal sulcus and transverse parietal sulci | 33 | -43 | 44 | 7.86 | 2594.9 | 0.0012 |
| Left angular gyrus | 33 | -65 | 45 | 7.41 |  |  |
| Left superior parietal lobule | 10 | -61 | 64 | 6.03 |  |  |
| Left precuneus | 14 | -75 | 46 | 5.94 |  |  |
| Left marginal branch of the cingulate sulcus | 16 | -39 | 42 | 10.96 | 1043.9 | 0.0072 |
| Right marginal branch of the cingulate sulcus | 13 | -28 | 38 | 7.7 | 2172.41 | 0.0008 |
| Right intraparietal sulcus and transverse parietal sulci | 22 | -63 | 43 | 6.34 |  |  |
| Right middle occipital gyrus | 40 | -80 | 30 | 5.99 |  |  |
| Right intraparietal sulcus and transverse parietal sulci | 36 | -46 | 36 | 5.99 | 941.89 | 0.0074 |
| Right supramarginal gyrus | 58 | -36 | 44 | 5.02 |  |  |
| Right calcarine sulcus | 12 | -75 | 6 | 4.18 | 457 | 0.0366 |
| Right calcarine sulcus | 25 | -55 | 1 | 3.9 |  |  |
| Right superior frontal gyrus | 7 | 0 | 59 | 5.13 | 450.49 | 0.037 |
| Right superior part of the precentral sulcus | 31 | -4 | 46 | 4.94 | 431.03 | 0.0406 |
| Right superior frontal gyrus | 18 | 14 | 62 | 4.66 |  |  |
| **Congenitally Blind Group > Adult-Onset Blind Group** |  |  |  |  |  |  |
| Right superior occipital gyrus | 14 | -92 | 15 | 4.67 | 483.51 | 0.046 |
| Right middle occipital gyrus | 30 | -89 | 12 | 4.53 |  |  |
| **Congenitally Blind Group > Sighted Group** |  |  |  |  |  |  |
| Left middle occipital gyrus | -34 | -88 | 14 | 4.98 | 528.72 | 0.0312 |
| Left middle occipital sulcus and lunatus sulcus | -25 | -95 | 1 | 4.78 |  |  |
| Right middle occipital sulcus and lunatus sulcus | 33 | -82 | 9 | 6.52 | 807.7 | 0.011 |
| Right medial occipito-temporal sulcus and lingual sulcus | 32 | -45 | -14 | 5.43 | 548.72 | 0.027 |

Brain regions active for language > math
|  | x | y | z | Peak t | mm2 | Pcluster |
| --- | --- | --- | --- | --- | --- | --- |
| **Adult-Onset Blind Group** |  |  |  |  |  |  |
| Left superior temporal gyrus | -61 | -15 | 3 | 11.81 | 3684.03 | 0.0002 |
| Left planum polare of the superior temporal gyrus | -47 | 7 | -17 | 11.14 |  |  |
| Left superior temporal sulcus | -51 | -49 | 5 | 9.74 |  |  |
| Left superior temporal sulcus | -54 | -19 | -15 | 8.88 |  |  |
| Left superior temporal sulcus | -41 | -63 | 19 | 7.08 |  |  |
| Left opercular part of the inferior frontal gyrus | -52 | 25 | 17 | 7.36 | 835.81 | 0.0242 |
| Left orbital sulci | -38 | 31 | -13 | 6.54 |  |  |
| Left precuneus | -5 | -61 | 31 | 7.28 | 781.82 | 0.0268 |
| Right superior temporal sulcus | 57 | -9 | -20 | 9.98 | 1576.82 | 0.0014 |
| Right lateral aspect of the superior temporal gyrus | 48 | 15 | -21 | 9.88 |  |  |
| Right lateral aspect of the superior temporal gyrus | 64 | -5 | -4 | 7.88 |  |  |
| **Congenitally Blind Group** |  |  |  |  |  |  |
| Left lateral aspect of the superior temporal gyrus | -61 | -14 | -5 | 13.01 | 4536.4 | 0.0002 |
| Left superior temporal sulcus | -53 | -39 | 3 | 9.03 |  |  |
| Left lateral aspect of the superior temporal gyrus | -46 | 16 | -26 | 8.68 |  |  |
| Left triangular part of the inferior frontal gyrus | -55 | 23 | 12 | 8.52 | 1166.59 | 0.0108 |
| Left orbital part of the inferior frontal gyrus | -47 | 32 | -14 | 7.08 |  |  |
| Left orbital gyri | -31 | 18 | -22 | 5.57 |  |  |
| Left superior frontal gyrus | -9 | 61 | 25 | 7.66 | 876.66 | 0.016 |
| Left subparietal sulcus | -10 | -55 | 26 | 9.64 | 799.73 | 0.019 |
| Right lateral aspect of the superior temporal gyrus | 65 | -10 | 0 | 12.51 | 5884.37 | 0.0002 |
| Right superior temporal sulcus | 49 | -13 | -15 | 12.07 |  |  |
| Right planum polare of the superior temporal gyrus | 39 | 9 | -27 | 9.7 |  |  |
| Right superior temporal sulcus | 51 | -60 | 19 | 9.41 |  |  |
| Right superior temporal sulcus | 45 | -40 | 3 | 8.55 |  |  |
| Right parahippocampal gyrus | 25 | -7 | -30 | 8.01 |  |  |
| Right anterior occipital sulcus and preoccipital notch | 45 | -69 | 10 | 7.09 |  |  |
| Right triangular part of the inferior frontal gyrus | 56 | 24 | 18 | 8.87 | 1309.03 | 0.0092 |
| Right triangular part of the inferior frontal gyrus | 52 | 32 | -4 | 7.72 |  |  |
| Right superior frontal gyrus | 10 | 56 | 32 | 7.64 | 978.34 | 0.0152 |
| Right superior frontal gyrus | 10 | 15 | 65 | 5.6 |  |  |
| Right lateral occipito-temporal gyrus (fusiform gyrus) | 38 | -48 | -22 | 8.69 | 891.01 | 0.017 |
| Right anterior transverse collateral sulcus | 41 | -8 | -35 | 6.59 |  |  |
| Right subparietal sulcus | 8 | -56 | 36 | 9.01 | 635.66 | 0.0278 |
| Right straight gyrus | 6 | 54 | -13 | 8.58 | 633.27 | 0.0282 |
| **Sighted Group** |  |  |  |  |  |  |
| Left lateral aspect of the superior temporal gyrus | -50 | 13 | -21 | 12.91 | 4568.42 | 0.0002 |
| Left superior temporal sulcus | -54 | -46 | 0 | 10.68 |  |  |
| Left lateral aspect of the superior temporal gyrus | -57 | -15 | -8 | 9.85 |  |  |
| Left superior temporal sulcus | -44 | -67 | 26 | 5.01 |  |  |
| Left horizontal ramus of the anterior segment of the lateral sulcus | -44 | 31 | -3 | 9.91 | 950.64 | 0.0112 |
| Left superior frontal gyrus | -6 | 55 | 32 | 10.72 | 902.06 | 0.0132 |
| Left superior frontal gyrus | -8 | 12 | 66 | 6.37 |  |  |
| Left subparietal sulcus | -12 | -51 | 36 | 8.66 | 811.73 | 0.0154 |
| Right lateral aspect of the superior temporal gyrus | 47 | 13 | -20 | 10.68 | 3109.97 | 0.0002 |
| Right lateral aspect of the superior temporal gyrus | 62 | -6 | -7 | 9.61 |  |  |
| Right superior temporal sulcus | 52 | -33 | 1 | 8.97 |  |  |
| Right middle temporal gyrus | 61 | -35 | -6 | 7.18 |  |  |
| Right precuneus | 5 | -58 | 31 | 6.23 | 642.25 | 0.018 |
| **Congenitally Blind Group > Adult-Onset Blind Group** |  |  |  |  |  |  |
| Right superior temporal sulcus | 51 | -6 | -17 | 6.01 | 2604.86 | 0.0002 |
| Right lateral occipito-temporal sulcus | 42 | -52 | -17 | 5.52 | 518.55 | 0.0436 |
| **Congenitally Blind Group > Sighted Group** |  |  |  |  |  |  |
| Right anterior occipital sulcus and preoccipital notch | 46 | -68 | 8 | 5.96 | 689.95 | 0.016 |
| Right lateral occipito-temporal sulcus | 42 | -50 | -18 | 6.14 | 533.68 | 0.0316 |
| Right calcarine sulcus | 17 | -74 | 9 | 5 | 477.16 | 0.0386 |

Peaks of brain regions active more for math than language ( $p < 0.05$ , cluster corrected; $p < 0.01$ cluster-forming threshold; 20 mm minimum distance between peaks). Coordinates reported in MNI space. Peak t: t values corresponding to local maxima; mm2: area occupied by cluster on cortical surface; Pcluster: P value for entire cluster

**Supplementary Table 3.**
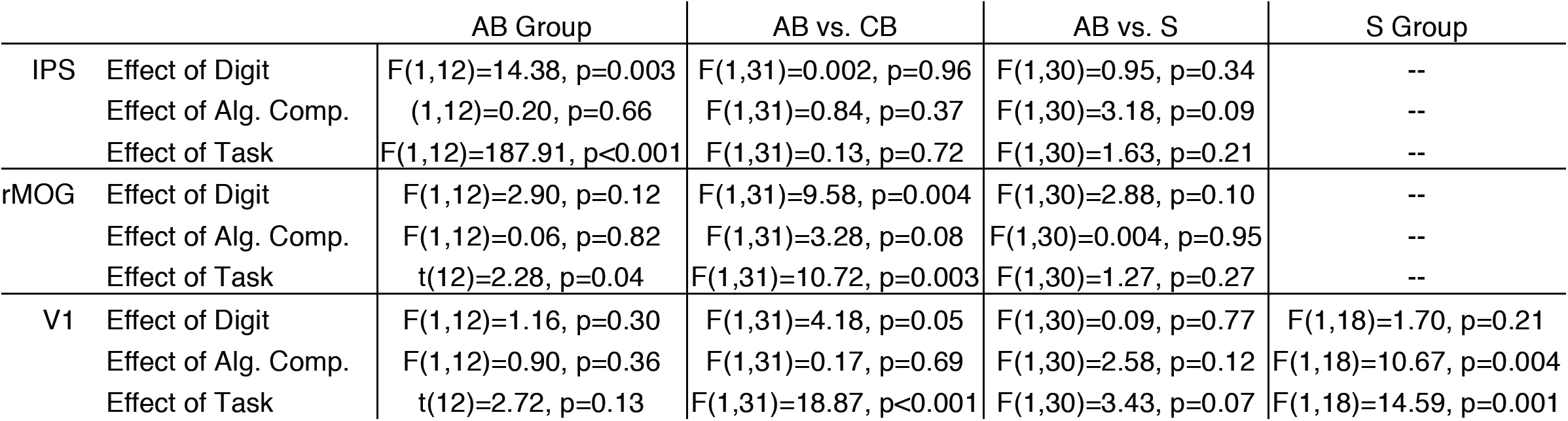
Summary of results from ROI analyses of math task.

|  |  | AB Group | AB vs. CB | AB vs. S | S Group |
| --- | --- | --- | --- | --- | --- |
| IPS | Effect of Digit | $F(1,12)=14.38, p=0.003$ | $F(1,31)=0.002, p=0.96$ | $F(1,30)=0.95, p=0.34$ | -- |
| | Effect of Alg. Comp. | $(1,12)=0.20, p=0.66$ | $F(1,31)=0.84, p=0.37$ | $F(1,30)=3.18, p=0.09$ | -- |
| | Effect of Task | $F(1,12)=187.91, p<0.001$ | $F(1,31)=0.13, p=0.72$ | $F(1,30)=1.63, p=0.21$ | -- |
| rMOG | Effect of Digit | $F(1,12)=2.90, p=0.12$ | $F(1,31)=9.58, p=0.004$ | $F(1,30)=2.88, p=0.10$ | -- |
| | Effect of Alg. Comp. | $F(1,12)=0.06, p=0.82$ | $F(1,31)=3.28, p=0.08$ | $F(1,30)=0.004, p=0.95$ | -- |
| | Effect of Task | $t(12)=2.28, p=0.04$ | $F(1,31)=10.72, p=0.003$ | $F(1,30)=1.27, p=0.27$ | -- |
| V1 | Effect of Digit | $F(1,12)=1.16, p=0.30$ | $F(1,31)=4.18, p=0.05$ | $F(1,30)=0.09, p=0.77$ | $F(1,18)=1.70, p=0.21$ |
| | Effect of Alg. Comp. | $F(1,12)=0.90, p=0.36$ | $F(1,31)=0.17, p=0.69$ | $F(1,30)=2.58, p=0.12$ | $F(1,18)=10.67, p=0.004$ |
| | Effect of Task | $t(12)=2.72, p=0.13$ | $F(1,31)=18.87, p<0.001$ | $F(1,30)=3.43, p=0.07$ | $F(1,18)=14.59, p=0.001$ |

**Supplementary Figure 1.**
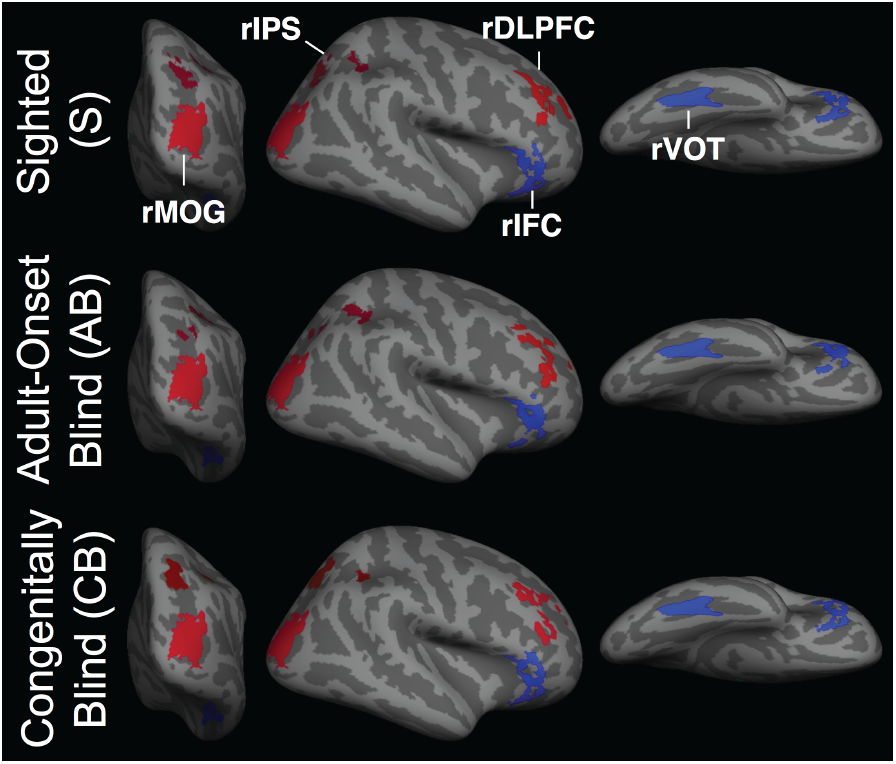
Resting-State Regions of Interest. Regions of interest (ROIs) used for resting-state analysis in sighted, adult-onset blind and congenitally blind groups. Red colors indicate ROIs of math network and blue colors indicate ROIs of language network. Visual cortex ROIs are identical across groups. Prefrontal and parietal ROIs defined separately for each group.

